# Putting the F in FBD analyses: tree constraints or morphological data ?

**DOI:** 10.1101/2022.07.07.499091

**Authors:** Joëlle Barido-Sottani, Alexander Pohle, Kenneth De Baets, Duncan Murdock, Rachel C. M. Warnock

## Abstract

The fossilized birth-death (FBD) process provides an ideal model for inferring phylogenies from both extant and fossil taxa. Using this approach, fossils (with or without character data) are directly considered as part of the tree. This leads to a statistically coherent prior on divergence times, where the variance associated with node ages reflects uncertainty in the placement of fossil taxa in the phylogeny. Since fossils are typically not associated with molecular sequences, additional information is required to place fossils in the tree. Previously, this information has been provided in two different forms: using topological constraints, where the user specifies monophyletic clades based on established taxonomy, or so-called total-evidence analyses, which use a morphological data matrix with data for both fossil and extant specimens in addition to the molecular alignment. In this work, we use simulations to evaluate these different approaches to handling fossil placement in FBD analyses, both in ideal conditions and in datasets including uncertainty or even errors. We also explore how rate variation in fossil recovery or diversification rates impacts these approaches. We find that the extant topology is well recovered under all methods of fossil placement. Divergence times are similarly well recovered across all methods, with the exception of constraints which contain errors. These results are consistent with expectations: in FBD inferences, divergence times are mostly informed by fossil ages, so variations in the position of fossils strongly impact these estimates. On the other hand, the placement of extant taxa in the phylogeny is driven primarily by the molecular alignment. We see similar patterns in datasets which include rate variation, however one notable difference is that relative errors in extant divergence times increase when more variation is included in the dataset, for all approaches using topological constraints, and particularly for constraints with errors. Finally, we show that trees recovered under the FBD model are more accurate than those estimated using non-FBD (i.e., non-time calibrated) inference. This result holds even with the use of erroneous fossil constraints and model misspecification under the FBD. Overall, our results underscore the importance of core taxonomic research, including morphological data collection and species descriptions, irrespective of the approach to handling phylogenetic uncertainty using the FBD process.

## 2 Introduction

Time-calibrated trees provide a crucial basis for hypothesis-testing in the life and earth sciences. Phylogenetic dating combines molecular and fossil evidence, allowing us to reconstruct a timeline of events that are otherwise not directly observable. Within a Bayesian framework, temporal evidence is incorporated via the tree prior or tree model. The fossilised birth-death (FBD) process explicitly combines the lineage diversification and fossil recovery processes, providing an ideal model for inferring phylogenies from both extant species and fossil specimens (Stadler, 2010). Using this approach, fossils (with or without character data) are directly considered as part of the tree (Heath et al., 2014; Gavryushkina et al., 2014, 2017; Zhang et al., 2016). This leads to a statistically coherent prior on divergence times, where the variance associated with node ages reflects the incompleteness of the fossil record, as well as uncertainty associated with the placement of fossil taxa in the phylogeny. Bayesian inference using the FBD process as a tree prior also allows for reliable estimation of the diversification and sampling parameters. This model has been successfully applied to datasets of living and fossil taxa (Schuster et al., 2018; Šmíd and Tolley, 2019; Thomas et al., 2020; Pohle et al., 2022).

The initial implementation of the FBD model assumed constant diversification (birth and death) and fossil recovery rates through the entire phylogeny (Heath et al., 2014). However, a wide range of factors, from biological and geological processes to collection practices, contribute to the probability a given organism will be sampled in the fossil record (Kidwell and Holland, 2002; Smith and McGowan, 2011; Benson et al., 2021; Raja et al., 2022). As a consequence, fossil recovery potential varies substantially across time, space and taxa. Other variables, such as environmental conditions and phenotypic traits, contribute to variation in diversification (speciation and extinction) rates. Extensions of the FBD process have integrated these variations into the model (Gavryushkina et al., 2014; Zhang et al., 2016), however, many empirical analyses are still done under the constant-rate assumption, for several reasons such as computational cost, lack of precise knowledge of rate changes, and ease of setup and interpretation.

One challenge in integrating fossil specimens in FBD analyses is that unlike extant species, fossils are only exceptionally associated with molecular sequences. As a result, additional information needs to be added to the inference to allow fossils to be placed in the tree topology. In previous research, this information has been provided in two different forms, which can be used separately or in combination. The first is topological constraints, which are constraints added by the user specifying that certain subsets of tips, extant or extinct, need to be monophyletic clades in the inferred phylogeny. These constraints are generally based on the taxonomy, with the constrained clades corresponding to genera and/or higher classifications. Topological constraints use information which is usually already available, and do not add computational complexity or cost to the inference. However, they do not easily accommodate uncertainty in the taxonomy, which is present even for well-known crown groups (Marx et al., 2016). Another approach is so-called total-evidence analyses (Ronquist et al., 2012), which uses a morphological data matrix, with data for both fossil and extant specimens, in addition to the molecular alignment. The morphological matrix is added to the inference along with a morphological substitution model and a morphological clock model, and contributes to the phylogenetic likelihood. Although total-evidence approaches better account for the underlying empirical data, they are more costly both in the time and effort required to assemble the matrix and in the added computational cost of the inference. In addition, the accuracy and precision of the inference has been found to be strongly dependent on the size of the morphological matrix (Barido-Sottani et al., 2020). Although these two approaches can in theory be combined in the same analysis, in practice they are often viewed as separate alternatives (for instance, Šmíd and Tolley (2019) uses constraints, while Thomas et al. (2020) uses a total-evidence approach).

In this work, we compare and evaluate different approaches to add fossil placement information in FBD analyses, both in ideal conditions and in datasets including uncertainty or even errors. We also use datasets containing variation in fossil recovery rates or speciation rates to explore whether ignoring rate variations impacts fossil placement approaches.

## 3 Methods

### 3.1 Simulations

#### 3.1.1 Trees and fossils

Our goal was to assess the performance of the FBD process, examining the impact of phylogenetic uncertainty and model misspecification. The parameters of the simulation were constrained to reflect values obtained for marine invertebrates.

Trees and fossils were simulated using the R packages TreeSim and FossilSim. The simulations were conditioned on the number of extant tips (*n* = 25), with constant speciation rate (*λ* = 0.11), extinction rate (*μ* = 0.1) and fossil recovery rate (*ψ* = 0.03). We assumed complete sampling at the present (i.e. the probability of extant species sampling *ρ* = 1). Parameters were selected such that the expected origin time was 250 Myr and the expected number of fossils was 100. Simulated data sets were filtered to select for trees with an origin time between 225 and 275 Myr and with between 80 and 120 fossils, and simulations which did not fit these two criteria were discarded.

We also simulated two sets of trees with (a) variable fossil recovery rates (*ψ*_1_ = 0.02 and *ψ*_2_ = 0.04), and (b) variable speciation rates (*λ*_1_ = 0.12 and *λ*_2_ = 0.06) and variable fossil recovery rates (*ψ*_1_ = 0.02 and *ψ*_2_ = 0.04). The speciation rate in set (a) and the extinction rate in sets (a) and (b) were fixed to the same values as in the constant birth-death simulation. In both variable sets, the variation in rates was linked to a trait that was simulated along each tree under a Brownian motion process. Traits were simulated with an initial value of 2.25 (for set (a)) or 1.5 (for set (b)) and variance = 0.01. Trait values were then assigned to two discrete types: values *<* 0 were assigned to type 1 (with rates *ψ*_1_ and *λ*_1_) and values *>* 0 were assigned to type 2 (with rates *ψ*_2_ and *λ*_2_). The final trees were kept if each trait value was assigned to at least 10% of the tips. The parameters of the BM process were calibrated so the final sampled trees contained an average of 8 to 11 trait changes.

We generated 50 replicate phylogenies for each of the three simulation conditions.

#### 3.1.2 Characters

Molecular sequence alignments of 1000 sites were simulated under the HKY +G model with five discrete gamma rate categories (*α* = 0.25). Branch rates were simulated under a lognormal uncorrelated clock model. For each tree replicate the average substitution rate was sampled from a gamma distribution with an expected value = 1, and shape and scale parameters = 2 and 0.5, respectively. The log of this rate was then used to define the mean of a lognormal distribution with variance = 0.01 from which branch specific rates were independently drawn.

Morphological data matrices of 50 or 300 characters were simulated under a binary state Mk model with a strict clock and a rate = 0.1.

### 3.2 Inference

We analysed each replicate using Bayesian phylogenetic inference in the BEAST2 framework (Bouckaert et al., 2019), under the constant rate FBD process implemented in the package Sampled Ancestors (SA) (Gavryushkina et al., 2014).

We examined the impact of five different ways of incorporating the phylogenetic uncertainty associated with fossils:

1. ossil samples without character data were assigned to the nearest extant ancestral node using clade constraints (designated as “correct constraints”)(Fig. 1C).
2. As in (1) but with 2% of fossils assigned to the wrong node, selected at random (designated as “constraints with errors”)(Fig. 1D).
3. Fossil samples without character data were assigned to the node above the nearest ancestral node (designated as “imprecise constraints”)(Fig. 1E).
4. As in (1) but 5 nodes picked at random in the tree were collapsed, meaning all constraints below that node were removed (designated as “collapsed constraints”)(Fig. 1F).
5. Character data (= 50 characters) was included for both fossil and extant samples, with no additional constraints (designated as “total-evidence with n=50”)(Fig. 1G).
6. Character data (= 300 characters) was included for both fossil and extant samples, with no additional constraints (designated as “total-evidence with n=300”)(Fig. 1G).

**Figure 1:**
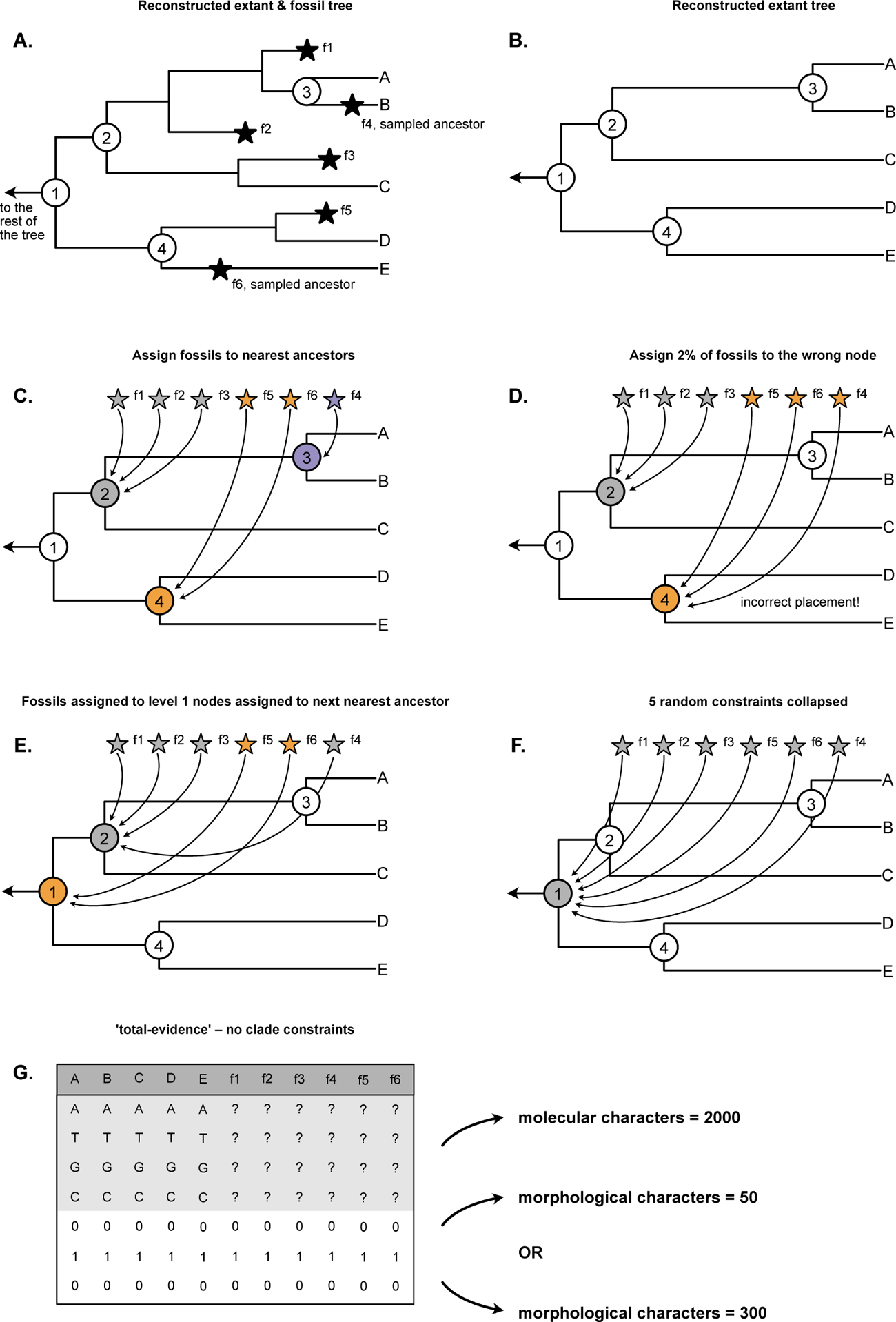
Schematic representation of the different analyses performed in this study. First, we simulate a full tree with fossil samples (A), from which the true extant tree can be obtained (B). We then set up clade constraints according to different rules: in (C), all clade constraints are correct and complete. In (D), some fossils are assigned to the wrong clade. In (E), all clades have a low level of taxonomic uncertainty. In (F), some subclades are fully unknown while the rest of the tree is fully known. Finally, we also perform two total-evidence analyses (G).

Configuration (1) corresponds to a situation with perfect information on fossil taxonomy (Fig. 1C), while configuration (2) includes a low amount of misplaced fossils (Fig. 1D). Configurations (3) and (4) include uncertainty in fossil placement in two different ways: in configuration (3) all clades have some amount of uncertainty in fossil assignments (Fig. 1E), in configuration (4) a few clades have no information on fossil placement within the clade, while the other clades are known perfectly (Fig. 1F). Configurations (5) and (6) are total-evidence analyses, with respectively a low or high amount of morphological characters for each fossil (Fig. 1G).

The FBD process was parameterized using the ‘canonical’ (speciation, extinction, sampling) parameterization, with an exponential prior (mean = 1.0) on the birth, death, sampling migration parameters. A uniform prior *U* (0, 1000) was used for the origin time of the process. We deliberately chose uninformative priors for these parameters in order to minimize the influence of the priors on the results. A lognormal prior for HKY parameter *κ* (mean = 1 and sd = 1.25), with a uniform prior on the state frequencies. An exponential prior for alpha for gamma distributed rates (mean = 1). The Mk model was applied to the binary character data. An exponential prior (mean = 1) was used for the mean of the uncorrelated relaxed clock model and a gamma prior (*α* = 0.5396 and *β* = 0.3819) on the standard deviation.

Analyses were run for a minimum of 200,000,000 generations, sampling every 10,000 generations and discarding 10% as burnin. We assessed convergence by calculating the ESS values for all model parameters – if any ESS values were *<* 200, analyses were longer (this was only necessary for 19 replicates). Eight analyses that failed to converge after 2,000,000,000 generations were excluded. We note that all of these are analyses in which fossils have no character data and are assigned to the next nearest ancestor (Fig. 1F). The maximum number excluded for a given simulation and inference scenario was 3.

For each replicate, we calculated the relative error of divergence time estimates (as the absolute difference between the median estimate and the true value, divided by the true value), averaged over all extant nodes, and the 95% HPD coverage, i.e. the proportion of extant nodes for which the true age was contained within the 95% Highest Posterior Density interval. We also measured the normalised Robinson-Foulds distance between the estimated phylogeny and the truth, averaged over all posterior samples, for both the full tree including fossils and the reconstructed extant phylogeny.

We also performed a separate unconstrained (i.e., non-FBD) tree inference using RevBayes. Since these analyses contain no age information the trees produced are non-time calibrated and have branch lengths in substitutions per site rather than in units of time.We used a uniform tree prior on the topology, with an exponential prior (*λ* = 10, mean = 0.1) on the branch lengths. We used the same settings as above for substitution models. We ran 4 independent chains for 200,000 generations, sampling every 200 generations, discarding 10% as burnin and combining the output. We assessed convergence by calculating the ESS values for all model parameters as above and examined the variance across chains. All runs converged. We measured the normalised Robinson-Foulds distance between the estimated phylogeny and the truth, averaged over all posterior samples, and compared those to the results obtained using the FBD inference.

## 4 Results

Figure 2 shows the results on the datasets simulated under constant rates, variable fossilization rates, and variable birth and fossilization rates.

**Figure 2:**
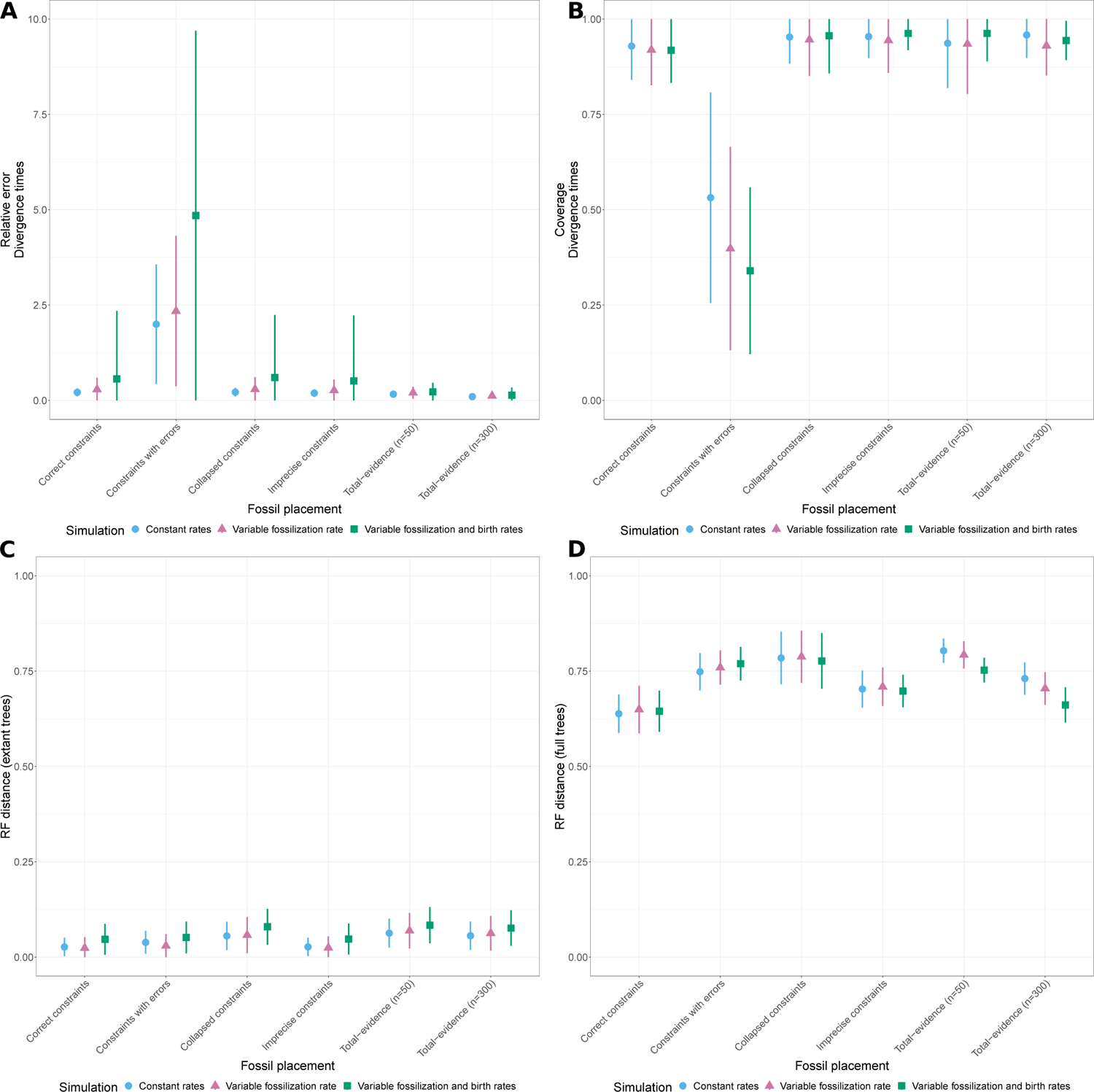
Results of FBD analyses for simulations under a constant birth-death process (red), variation in fossilization rates (green) or variation in birth and fossilization rates (blue). Absolute relative error (A) and 95% HPD coverage (B) of divergence time estimates, averaged over all nodes in the extant phylogeny. Average Robinson-Foulds distance between posterior samples and the true tree, for the extant tree only (C) or the full tree including fossil samples (D). All measures show the average and standard deviation across all replicates for different fossil placement approaches.

Under a constant-rate process, we can see that the extant topology is well recovered under all methods of fossil placement. Divergence times are similarly well recovered across all methods, with the marked exception of constraints which contain errors. Indeed, the relative error is much higher and the coverage much lower in the presence of errors. Overall, these results are quite consistent with expectations: in FBD inferences, divergence times are mostly informed by fossil ages, so variations in the position of fossils will strongly impact these estimates. On the other hand, the placement of extant taxa in the phylogeny is driven primarily by the molecular alignment, and thus is mostly independent from the handling of fossil specimens. As expected, fossil placement methods also impact the accuracy of the estimate of the full phylogeny. The best performance is obtained with correct constraints, followed by imprecise constraints and total-evidence with a high number of morphological characters, whereas the worst estimates are obtained using total-evidence with a low number of morphological characters.

We see very similar patterns in datasets which include rate variation. One notable difference is that relative errors in extant divergence times are higher the more variation is included in the dataset for all approaches using topological constraints, and particularly for constraints with errors. Interestingly, total-evidence approaches show the same levels of relative error in extant divergence times regardless of rate variation in the simulation models. One possible explanation is that total-evidence approaches include more uncertainty than topological constraints and so are able to better compensate for model mismatches between the simulation and the inference. The effect of simulated rate variation is almost null on the other metrics, i.e. the coverage of extant divergence times and the RF distances for extant and full phylogenies.

Figure. 3 shows the results obtained using the FBD versus non-FBD (i.e., without fossil age information) inference. The comparison shows that trees recovered under the FBD model are more accurate, in terms of Robinson-Foulds distance – this applies to both the extant topology estimated using the molecular data only or the total-evidence matrix and the full topology. This result holds even with the use of erroneous fossil constraints and model misspecification under the FBD. Interestingly, the unconstrained topology estimates are more accurate for trees simulated using constant diversification and sampling rates than trees simulated with rate variation. This might reflect a better correspondence between the uniform tree model and a constant birth-death sampling process. The inclusion of fossil taxa increases the accuracy of the extant tree. More morphological character data also leads to improved accuracy of both the extant and full trees.

**Figure 3:**
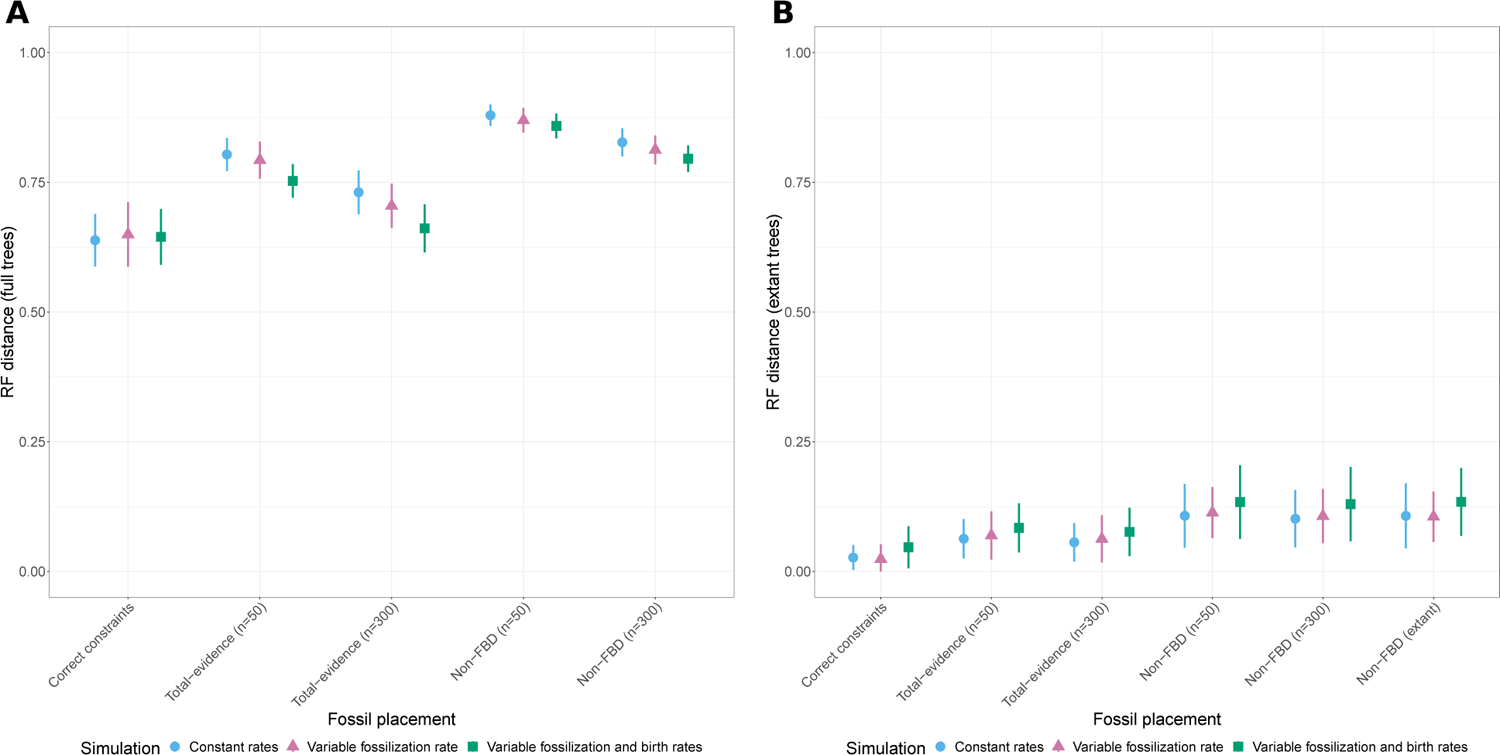
Comparison with non-FBD analysis (i.e., with no fossil age information) for simulations under a constant birth-death process (red), variation in fossilization rates (green) or variation in birth and fossilization rates (blue). Average Robinson-Foulds distance between posterior samples and the true tree, for the extant tree only (C) or the full tree including fossil samples (D). All measures show the average and standard deviation across all replicates for different fossil placement approaches.

## 5 Discussion

The fossilised birth-death process offers flexible opportunities to reconstruct dated phylogenies incorporating fossils under a mechanistic framework. Here, we explored different options for handling the phylogenetic or taxonomic uncertainty associated with fossil samples using simulations.

We found that alternative approaches to including fossils had relatively little impact on the accuracy of the extant topology (based on RF distances, Fig. 2C-D). As noted above, the signal for the extant topology largely comes from the molecular sequence alignment, which explains why the extant phylogeny was reasonable even when a small portion of fossils were placed erroneously in the tree. Using fixed constraints to assign fossils to nodes produced more accurate trees, compared to trees recovered using morphological characters to place the fossils. However, this represents an idealised scenario, as reliable and precise taxonomic information used to inform constraints will often not be available (see below for further discussion on data quality and availability). The results obtained using non-time calibrated tree inference show that trees recovered with the inclusion of fossils with character data are more accurate than trees based on the extant taxa only (Fig. 3A-B). We also confirm that results are even better when the fossil age information is taken into account using the FBD process. Previous simulations studies have also shown that the inclusion of fossils and age information can improve the accuracy of the topology among extant taxa (Mongiardino Koch et al., 2021).

Different approaches to handling taxonomic uncertainty had a more notable impact on divergence times. We found that the best possible scenario, in terms of recovering accurate node ages (relative error and coverage), was to assign fossils to the correct nearest node in the extant tree (Fig. 2A-B). This is unsurprising, as this is equivalent to fixing large portions of the topology using correct monophyletic clade constraints. But we further show that the use of less precise clade constraints (i.e., assigning fossils to larger or more inclusive clades, Fig. 1E) also recovers accurate estimates, similar to those obtained inferring the position of fossils based on morphological character data. Similarly, Heath et al. (2014) found using simulations that less precise constraints increased the variance but did not reduce accuracy of posterior divergence estimates (see also O’Reilly and Donoghue (2020)). However, our results show that when only a small proportion (2%) of fossils are assigned to incorrect nodes, overall accuracy decreases. Assigning larger proportions of fossils to incorrect nodes will reduce accuracy further. Preliminary runs using 10% of fossils assigned to incorrect nodes lead to coverage *<* 0.1 (results not shown). Together, our results suggest that uncertainty is a much less critical issue for inference than outright errors. Thus, we recommend that when in doubt, it is better to err on the side of caution and to either use larger and more inclusive clade constraints, since this does not compromise the overall accuracy of results and only leads to a small overall decrease in precision, or to infer the position of fossils using morphological data when available. The latter is preferential since the posterior will best reflect uncertainty associated with the placement of fossil taxa. In addition, if clade constraints are overly conservative (i.e., based on very large clades or higher level taxonomic divisions) there will be too much uncertainty and the analyses might fail to converge. ‘Imprecise constraints’ was the only simulation scenario in which we had to exclude replicates due to lack of convergence.

We also examined the impact of model violation on the accuracy of topology and divergence times recovered under the FBD model, by varying sampling rate only or both sampling and diversification rates. Not accounting for rate variation in these parameters has a modest impact on the accuracy of the topology *−* again reflecting the fact that the signal for topology predominantly comes from the character data (Fig. 2C-D). Model misspecification has a much more discernible impact on the divergence estimates (Fig. 2A-B). This is because the signal for divergence times largely comes from the distribution of fossil sampling times and relies more on the birth-death sampling model being correct. However, in most scenarios coverage for node ages remains high. These general results also match the findings of previous simulation studies that explored model misspecification under the FBD process (Heath et al., 2014) There are challenging identifiability issues associated with birth-death processes (Louca and Pennell, 2020; Louca et al., 2021). In particular, Louca and Pennell (2020) identified ‘congruence classes’, within which infinitely many diversification histories can have the same likelihood. This means that applying oversimplified birth-death models can result in highly misleading results. Our findings show that reliable phylogenies can be obtained under the FBD process, even if the underlying process is more complex than the model used for inference. The most notable exception are the results obtained with erroneous fossil constraints – when a portion of fossils are incorrectly placed in the tree, the relative negative impact of model misspecification is worse. Since we can rarely be certain of the underlying diversification or sampling process, this further underscores the need to use taxonomic constraints with extreme caution. Furthermore, our results demonstrate that total-evidence analyses are more robust against model misspecification, adding to the list of benefits associated with this approach.

We note that our simulation design represents an idealised scenario in several ways. For instance, we only simulated binary characters for our morphological alignments, whereas empirical data matrices often include multi-state characters or characters with hierarchical state dependencies. In addition, our matrices had no missing data, which is unrealistic for empirical datasets including fossils. These two factors likely contribute to the good overall performance of the total-evidence analyses in our study.

Ordinarily clade constraints and ‘total-evidence’ analyses are treated as two distinct approaches, but they can easily be combined. In fact this might be desirable, since several previous simulation studies have shown that the accuracy of parameters estimated under the FBD process increases with fossil sampling (Heath et al., 2014; Barido-Sottani et al., 2020). In some cases, we might only have abundant morphological character data associated with a subset of fossils. For example, we might have fossils with fewer traits (e.g., fossil cephalopod shells), alongside rarer but exceptionally preserved specimens with more informative traits (e.g., fossil cephalopods associated with jaws and or soft-tissues). Shell characteristics that are diagnostic of higher-level taxonomic groups could be used to define clade constraints for specimens associated with a smaller amount of data. Similarly, in palaeobotany, rarely preserved tissues (e.g., flowers or seeds) can have a small number of traits (i.e., 2-3) that would be considered too few to construct a matrix for phylogenetic analysis, but that are nonetheless considered definitive synapomorphies of certain plant groups. Another example could be ammonoid aptychi or lower jaws, which have been mapped to phylogenies based on a small number of specimens with known jaws, relative to those where shells/moulds are known (Engeser and Keupp, 2002).

We might also have different types of evidence associated with the presence of a group. For instance, worm eggs can sometimes be assigned to higher taxonomic ranks based on the presence of specific structures (Hugot et al., 2014), but these traits can have little to do with the adult morphology of exceptionally preserved worms. Other examples could include trace fossils (e.g., where they predate body fossils), molecular fossils (e.g., biomarkers), or exceptional preservation of different ontogenetic stages (e.g., small larvae in early phosphatic or Orsten-type preservation, see Maas et al. (2006)). In this way, we can partition different types of evidence to use in different ways. Parasites might be an especially good example in this context. Many groups of parasites are highly diverse today but most have poor preservation in the past, associated with two distinct fossil datasets *−* rare exceptionally preserved specimens that can be used to build a morphological matrix versus more common eggs or trace fossils that can only be assigned to higher groupings (De Baets et al., 2021), potentially better suited to the use of clade constraints (Warnock and Engelstädter, 2021).

Here, we considered a scenario in which we have an extant clade. For fossil only datasets, typically we use the FBD model with morphological data only and no topological constraints. Previous simulations have shown that trees generated with matrices that are typical of many fossil groups (i.e., 30 characters) will be highly uncertain (Barido-Sottani et al., 2020). Clade constraints could also be useful in this context, provided these can be defined with a high degree of confidence. As an example, a backbone constraint based on the analysis of molecular data could be applied to a tree that consists mostly fossils or is based on morphology, but where the morphology never recovers the established molecular phylogeny. For example, cyclostomes or many mammalian superorders are never recovered as monophyletic on the basis of morphology alone. In other cases, we might want to look at a large clade that contains too many fossil species for all of them to be included in a single analyses, but where previous studies show unequivocal support for certain subclades. In this scenario, clade constraints could help stabilising the topology, without the need to collect additional morphological data while remaining computationally efficient.

For some groups (e.g., soft-bodied worms or unicellular organisms), we it might not be possible or practical to collect much more data, due to both incomplete preservation and/or the labour associated with collecting morphological data, or obtain reliable information about broader level taxonomy. In this situation, extended versions of the FBD process can help make the best use of the available data. For instance, the occurrence birth-death process allows fossils associated with morphological character data and fossils associated with age data only to be assigned different sampling rates. Under this model, occurrence data do not need to be constrained to any part of the tree, but can improve overall FBD parameter estimates.

Beyond inferring dated trees, Soul and Friedman (2015) showed that trees constructed using higher level taxonomy and dated using time-scaling methods can be successfully used for phylogenetic comparative analyses. Although we emphasise the need to take a very cautious approach when using clade constraints, the FBD model could also be used in this context, with the added benefit that the output better reflects uncertainty in fossil ages, node ages and phylogenetic uncertainty.

Irrespective of the approach to handling taxonomic or phylogenetic uncertainty using the FBD process, core taxonomic research, including morphological data collection and species descriptions, remain essential. The issues we observed with the erroneous placement of fossils mirror those identified previously in node dating studies, where node calibrations based on inaccurate fossil assignments have been shown to result in large errors in divergence time estimation (e.g. Phillips et al., 2009). These issues spurred the development of ‘best practices’ for justifying fossil calibrations for node dating, taking into account both phylogenetic and fossil age uncertainty (Parham et al., 2012). Authors are recommended to provide an explicit set of statements justifying the assignment of a fossil to a given node, with reference to up-to-date phylogenetic analyses incorporating the relevant taxa or to a set of unequivocal synapomorphies. This rigorous and transparent approach to defining clade constraints is directly applicable to analyses that employ the FBD process, along with the criteria used to justify fossil ages. How age uncertainty is handled in analyses using the FBD process also has important implications for the accuracy of both divergence times and topology (Barido-Sottani et al., 2019, 2020). Developments of phylogenetic models used in palaeobiology are no substitute for the expertise contributed by fundamental systematics, taxonomic and stratigraphic research. Our results reiterate the need for increased and direct support for taxonomy-based projects (Britz et al., 2020; Engel et al., 2021). Improving approaches to phylogenetic dating requires both advanced methodological and empirical perspectives.

## 6 Acknowledgments

JBS was supported by funds from the National Science Foundation (USA), grants DBI-1759909 and DEB-1556615 and from the European Union’s Horizon 2020 Research and Innovation Programme under the Marie Sklodowska-Curie grant agreement No. 101022928. KDB was supported by the I.3.4 Action of the Excellence Initiative *−* Research University Programme at the University of Warsaw (Project: PARADIVE).

## References

Jöelle Barido-Sottani, Gabriel Aguirre-Fernández, Melanie J Hopkins, Tanja Stadler, and Rachel Warnock. Ignoring stratigraphic age uncertainty leads to erroneous estimates of species divergence times under the fossilized birth–death process. Proceedings of the Royal Society B, 286 (1902):20190685, 2019.

Jöelle Barido-Sottani, Nina van Tiel, Melanie J Hopkins, David F Wright, Tanja Stadler, and Rachel Warnock. Ignoring fossil age uncertainty leads to inaccurate topology and divergence time estimates in time calibrated tree inference. Frontiers in Ecology and Evolution, 8:183, 2020.

Roger BJ Benson, Richard Butler, Roger A Close, Erin Saupe, and Daniel L Rabosky. Biodiversity across space and time in the fossil record. Current Biology, 31(19):R1225–R1236, 2021.

Remco Bouckaert, Timothy G. Vaughan, Jöelle Barido-Sottani, Sebastián Duchêne, Mathieu Fourment, Alexandra Gavryushkina, Joseph Heled, Graham Jones, Denise Kühnert, Nicola De Maio, Michael Matschiner, Fábio K. Mendes, Nicola F. Müller, Huw A. Ogilvie, Louis Du Plessis, Alex Popinga, Andrew Rambaut, David Rasmussen, Igor Siveroni, Marc A. Suchard, Chieh Hsi Wu, Dong Xie, Chi Zhang, Tanja Stadler, and Alexei J. Drummond. BEAST 2.5: An advanced software platform for Bayesian evolutionary analysis. PLoS Computational Biology, 15(4):e1006650, 2019. ISSN 15537358. doi: 10.1371/journal.pcbi.1006650.

Ralf Britz, Anna Hundsdörfer, and Uwe Fritz. Funding, training, permits—the three big challenges of taxonomy. Megataxa, 1(1):49–52, 2020.

Kenneth De Baets, John Warren Huntley, Adiël A. Klompmaker, James D. Schiffbauer, and A. D. Muscente. The fossil record of parasitism: Its extent and taphonomic constraints. In Kenneth De Baets and John Warren Huntley, editors, The Evolution and Fossil Record of Parasitism: Coevolution and Paleoparasitological Techniques, pages 1–50. Springer International Publishing, Cham, 2021. ISBN 978-3-030-52233-9. doi: 10.1007/978-3-030-52233-9_1.

Michael S Engel, Luis MP Ceríaco, Gimo M Daniel, Pablo M Dellapé, Ivan Löbl, Milen Marinov, Roberto E Reis, Mark T Young, Alain Dubois, Ishan Agarwal, et al. The taxonomic impediment: a shortage of taxonomists, not the lack of technical approaches. Zoological Journal of the Linnean Society, 193(2):381–387, 2021.

Theo Engeser and Helmut Keupp. Phylogeny of the aptychi-possessing neoammonoidea (apty-chophora nov., cephalopoda). Lethaia, 35(1):79–96, 2002.

Alexandra Gavryushkina, David Welch, Tanja Stadler, and Alexei J Drummond. Bayesian inference of sampled ancestor trees for epidemiology and fossil calibration. PLoS Computational Biology, 10(12):e1003919, 2014.

Alexandra Gavryushkina, Tracy A Heath, Daniel T Ksepka, Tanja Stadler, David Welch, and Alexei J Drummond. Bayesian total-evidence dating reveals the recent crown radiation of penguins. Systematic biology, 66(1):57–73, 2017.

Tracy A Heath, John P Huelsenbeck, and Tanja Stadler. The fossilized birth–death process for coherent calibration of divergence-time estimates. Proceedings of the National Academy of Sciences, 111(29):E2957–E2966, 2014.

Jean-Pierre Hugot, Scott L Gardner, Victor Borba, Priscilla Araujo, Daniela Leles, Átila Augusto Stock Da-Rosa, Juliana Dutra, Luiz Fernando Ferreira, and Adauto Araújo. Discovery of a 240 million year old nematode parasite egg in a cynodont coprolite sheds light on the early origin of pinworms in vertebrates. Parasites & Vectors, 7(1):486, 2014. ISSN 1756-3305. doi: 10.1186/s13071-014-0486-6. URL https://doi.org/10.1186/s13071-014-0486-6.

Susan Kidwell and Steven Holland. The quality of the fossil record: Implications for evolutionary analyses. Annual Review of Ecology and Systematics, 11:561–88, 11 2002. doi: 10.1146/annurev.ecolsys.33.030602.152151.

Stilianos Louca and Matthew W Pennell. Extant timetrees are consistent with a myriad of diver-sification histories. Nature, 580(7804):502–505, 2020.

Stilianos Louca, Angela McLaughlin, Ailene MacPherson, Jeffrey B Joy, and Matthew W Pennell. Fundamental identifiability limits in molecular epidemiology. Molecular biology and evolution, 38(9):4010–4024, 2021.

Andreas Maas, Andreas Braun, Xi-Ping Dong, Philip C.J. Donoghue, Klaus J. Müller, Ewa Olempska, John E. Repetski, David J. Siveter, Martin Stein, and Dieter Waloszek. The ‘orsten’—more than a cambrian konservat-lagerstätte yielding exceptional preservation. Palaeoworld, 15(3): 266–282, 2006. ISSN 1871-174X. doi: https://doi.org/10.1016/j.palwor.2006.10.005. URL https://www.sciencedirect.com/science/article/pii/S1871174X06000382. The Fourth International Symposium on the Cambrian System.

Felix Marx, Olivier Lambert, and Mark D Uhen. Cetacean Paleobiology. Wiley-Blackwell, 05 2016. ISBN 978-1-118-56127-0.

Nicolás Mongiardino Koch, Russell J Garwood, and Luke A Parry. Fossils improve phylogenetic analyses of morphological characters. Proceedings of the Royal Society B, 288(1950):20210044, 2021.

Joseph E O’Reilly and Philip CJ Donoghue. The effect of fossil sampling on the estimation of divergence times with the fossilized birth–death process. Systematic biology, 69(1):124–138, 2020.

James F Parham, Philip CJ Donoghue, Christopher J Bell, Tyler D Calway, Jason J Head, Patricia A Holroyd, Jun G Inoue, Randall B Irmis, Walter G Joyce, Daniel T Ksepka, et al. Best practices for justifying fossil calibrations. Systematic Biology, 61(2):346–359, 2012.

Matthew J Phillips, Thomas H Bennett, and Michael SY Lee. Molecules, morphology, and ecology indicate a recent, amphibious ancestry for echidnas. Proceedings of the National Academy of Sciences, 106(40):17089–17094, 2009.

Alexander Pohle, Björn Kröger, Rachel Warnock, Andy H King, David H Evans, Martina Aubrechtová, Marcela Cichowolski, Xiang Fang, and Christian Klug. Early cephalopod evolution clarified through bayesian phylogenetic inference. BMC biology, 20(1):1–30, 2022.

Nussaïbah B Raja, Emma M Dunne, Aviwe Matiwane, Tasnuva Ming Khan, Paulina S Nätscher, Aline M Ghilardi, and Devapriya Chattopadhyay. Colonial history and global economics distort our understanding of deep-time biodiversity. Nature ecology & evolution, 6(2):145–154, 2022.

F. Ronquist, S. Klopfstein, L. Vilhelmsen, S. Schulmeister, D. L. Murray, and A. P. Rasnitsyn. A total-evidence approach to dating with fossils, applied to the early radiation of the Hymenoptera. Systematic Biology, 61:973–999, 2012.

Astrid Schuster, Sergio Vargas, Ingrid S Knapp, Shirley A Pomponi, Robert J Toonen, Dirk Erpenbeck, and Gert Wörheide. Divergence times in demosponges (Porifera): first insights from new mitogenomes and the inclusion of fossils in a birth-death clock model. BMC Evolutionary Biology, 18(1):1–11, 2018.

Jiří Šmíd and Krystal A Tolley. Calibrating the tree of vipers under the fossilized birth-death model. Scientific Reports, 9(1):1–10, 2019.

Andrew B. Smith and Alistair J. McGowan. The ties linking rock and fossil records and why they are important for palaeobiodiversity studies. Geological Society, London, Special Publications, 358(1):1–7, 2011. ISSN 0305-8719. doi: 10.1144/SP358.1. URL https://sp.lyellcollection.org/content/358/1/1.

Laura C Soul and Matt Friedman. Taxonomy and phylogeny can yield comparable results in comparative paleontological analyses. Systematic Biology, 64(4):608–620, 2015.

Tanja Stadler. Sampling-through-time in birth–death trees. Journal of Theoretical Biology, 267(3): 396–404, 2010.

Daniel B. Thomas, Alan J. D. Tennyson, R. Paul Scofield, Tracy A. Heath, Walker Pett, and Daniel T. Ksepka. Ancient crested penguin constrains timing of recruitment into seabird hotspot. Proceedings of the Royal Society B: Biological Sciences, 287(1932):20201497, 2020. doi: 10.1098/rspb.2020.1497. URL https://royalsocietypublishing.org/doi/abs/10.1098/rspb.2020.1497.

Rachel Warnock and Jan Engelstädter. The molecular clock as a tool for understanding host-parasite evolution. In Kenneth De Baets and John Warren Huntley, editors, The Evolution and Fossil Record of Parasitism: Coevolution and Paleoparasitological Techniques, pages 417–450. Springer International Publishing, Cham, 2021. ISBN 978-3-030-52233-9. doi: 10.1007/978-3-030-52233-9_13.

Chi Zhang, Tanja Stadler, Seraina Klopfstein, Tracy A Heath, and Fredrik Ronquist. Total-evidence dating under the fossilized birth–death process. Systematic Biology, 65(2):228–249, 2016.

